# Respiration facilitates behaviour during multisensory integration

**DOI:** 10.1101/2025.01.10.632352

**Authors:** Martina Saltafossi, Andrea Zaccaro, Daniel S. Kluger, Mauro Gianni Perrucci, Francesca Ferri, Marcello Costantini

## Abstract

The brain processes information from the external environment alongside signals generated by the body. Among bodily rhythms, respiration emerges as a key modulator of sensory processing. Multisensory integration, the non-linear combination of information from multiple senses to reduce environmental uncertainty, may be influenced by respiratory dynamics. This study investigated how respiration modulates reaction times and multisensory integration in a simple detection task. Forty healthy participants were presented with unimodal (Auditory, Visual, Tactile) and bimodal (Audio-Tactile, Audio-Visual, Visuo-Tactile) stimuli while their respiratory activity was recorded. Results revealed that reaction times systematically varied with respiration, with faster responses during peak inspiration and early expiration but slower responses during the expiration-to-inspiration transition. Applying the race model inequality approach to quantify multisensory integration, we found that Audio-Tactile and Audio-Visual stimuli exhibited the highest integration during the expiration-to-inspiration phase. These findings conceivably reflect respiration phase-locked changes in cortical excitability which in turn, orchestrates multisensory integration. Interestingly, participants also tended to adapt their respiratory cycles, aligning response onsets preferentially with early expiration. This suggests that, rather than a mere bottom-up mechanism, respiration is actively adjusted to maximise the signal-to-noise balance between interoceptive and exteroceptive signals.

## Introduction

The brain dynamically interacts with the external world, processing information from the five senses and guiding behavioural responses. Current research attempts to recast neural implementation of cognition to include the modulatory effects of peripheral bodily signals, i.e., interoception [1–3]. Human respiration represents a unique interoceptive system because, besides being a simple reflex, it can also be under volitional control [4–6]. In addition, breathing patterns are continuously adjusted to match not only metabolic demands, but also vocal behaviours (e.g., speaking, crying, laughing and singing) as well as vital functions (e.g., swallowing and suckling) [4,7,8]. Recent animal and human studies revealed a profound intertwining between the respiratory cycle and spontaneous brain activity, extending beyond homeostatic and allostatic needs [9]. Through multiple, and not necessarily independent, olfactory, somatosensory, interoceptive, and chemoreceptive pathways and mechanisms [10], respiration drives changes in the amplitude of brain oscillations [11–17], shifts of cortical and cortico-spinal excitability [18–20], and modulations of neuronal spiking in cortical and subcortical structures [15,21,22].

As a functional consequence of this respiratory entrainment of neural dynamics, cycle-to-cycle coupling of breathing and behaviour has been repeatedly found [23–26]. For instance, inspiration has been shown to improve recognition of fearful faces [27], memory of odours [28], accuracy in visuospatial [11,29] and perceptual decision making tasks [30], as well as to reduce reaction times (RTs) to visual [31] and auditory stimuli [32]. A similar phase dependency has been revealed for interoceptive attention optimisation [33,34], startle responses to auditory inputs [35], conscious tactile perception [36], conditioned learning [37,38], and action execution [39,40] all being boosted by expiration. From a predictive processing perspective, which posits that perception results from minimisation of prediction errors (PEs) (i.e. the discrepancy between priors and sensory input) [41], such covariation between respiration and behaviour has been interpreted as an *active sensing* mechanism [42]. This hypothesis rests on animal studies demonstrating that rodents’ sensory-motor routines (e.g., orofacial movements like sniffing and whisking) are gated to the respiratory rhythm [43–45]. Similarly, human participants spontaneously adapt their respiratory cycle to the onset of stimuli and responses, preferentially matching inspiration or expiration, likely depending on task difficulty and sensory modality [23,27,29,36,46]. By aligning events with respiration, the brain might fine-tune neural processes underlying adaptive behaviour [9,10,39].

Overall, the research examined so far highlights the crucial role of respiration in regulating information gating and interaction with the environment, by acting as a “clock mechanism” [11,43,45]. However, studies on respiratory modulations of perception have largely investigated one single sensory modality at a time. Hence, it remains unknown whether and how respiration modulates perception of everyday multisensory stimuli. Here, each conscious percept stems from the continuous combination and integration of multiple sensory inputs from the external world, which results in enhanced neural and behavioural outputs [47–50]. This facilitation provided by the synthesis among multiple sources of information is defined as multisensory integration (MI) and is experimentally assessed by comparing responses to cross-modal stimuli with those of the corresponding unisensory stimuli (i.e., effectiveness) [51–53].

We recently demonstrated a significant interplay between exteroceptive (multisensory) and interoceptive processes by showing that the cardiac cycle modulates multisensory integration, likely through competition-like suppression of tactile stimuli during systole [54]. Here, we extended this investigation to explore the coupling between respiration and multisensory perception. We subjected participants to a simple detection task involving three sensory modalities (i.e., hearing, touch, and vision), presented alone (unimodal) or in combination (bimodal). We first hypothesised that speeded responses to unimodal and bimodal inputs would vary systematically across phases of spontaneous respiration. Second, after assessing super-additive interactions during bimodal stimulations, we tested whether the magnitude of multisensory integration was modulated across four respiratory phase-bins, segmenting the cycle into transition (inspiration-to-expiration and expiration-to-inspiration) and non-transition (inspiration and expiration) phases. On the behavioural level, we examined the alignment of response onsets to respiration phase in keeping with previous reports of sensory-motor coupling (see above).

## Materials and methods

### Participants

Forty-one participants (29 female; 3 left-handed; mean age ± SD = 24.88 ± 2.90 years) took part in the study, recruited from the “Gabriele d’Annunzio” University of Chieti-Pescara and the wider community. All participants had normal or corrected-to-normal vision. Exclusion criteria for participating were self-reported history of hearing loss and either mental, cardiovascular, or neurological disorders. Before the experiment, participants gave written informed consent. Ethical approval from the local ethics board was obtained (Institutional Review Board of Psychology of the Department of Psychological, Health and Territorial Sciences, “Gabriele d’Annunzio” University of Chieti - Pescara, Protocol Number 23013). The experiment was conducted following the Declaration of Helsinki.

Analyses were performed on 40 participants after excluding one participant due to excessive missed responses (> 50%). Block 1 was excluded from 2 participants due to failures in data recordings. A total of 28,320 trials were analysed.

### Experimental setup

The stimulus delivery apparatus was identical to that described in [54] (**Figure 1a**). The stimulation included three unimodal stimuli—Auditory (A), Tactile (T), and Visual (V)—as well as their bimodal combinations: Audio-Tactile (AT), Audio-Visual (AV), and Visuo-Tactile (VT). For a detailed explanation of the stimulus types and thresholding procedure, readers are referred to Supplementary 1 and Saltafossi et al. (2023). Respiratory activity was recorded with a BIOPAC MP160 (BIOPAC System, Inc., Goleta, CA, USA) (Low-pass filter: 35 Hz; high-pass filter: 0.05 Hz; notch filter: 50 Hz; sampling rate: 2000 Hz) using AcqKnowledge software (version 5.0.5, BIOPAC System, Inc., Goleta, CA, USA). A respiration belt with a transducer was placed around the participants’ chest.

**Figure 1.**
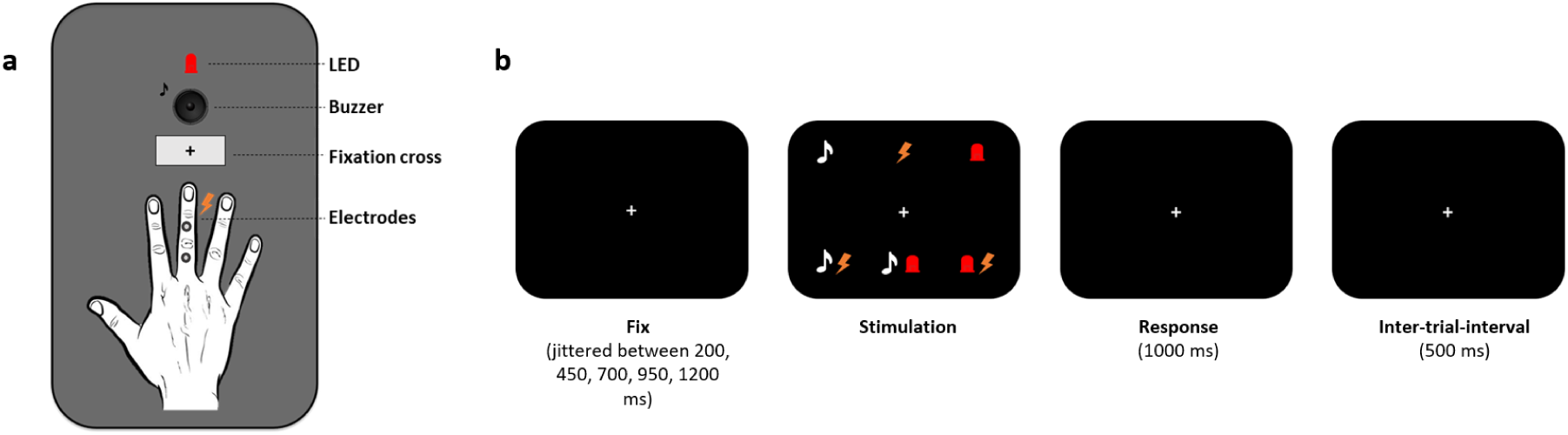
Experimental setup. (**a**) Stimulus delivery apparatus: the in-house box presented electric pulses as tactile stimuli, brief flashes as visual stimuli (LED), and auditory stimuli via a buzzer, in close proximity. (**b**) Timeline of trials: after a period with a variable duration, a stimulation occurred (either unimodal or bimodal). Participants had to respond within 1000 msec by pressing a pedal. The response window was then followed by an inter-trial interval of 500 msec.

The experiment design mirrored that of Saltafossi et al. (2023) but stimuli were not triggered by cardiac or respiratory phases. In a simple detection task, participants fixated on a central cross and responded quickly to stimuli. After a variable delay (200–1200 ms), unimodal (A, T, V) or bimodal (A, AV, VT) stimuli were presented. Participants pressed a pedal with their right foot upon perceiving a stimulus, with a maximum response time of 1000 ms. Each trial had a fixed 500 ms inter-trial interval (**Figure 1b**). A total of 720 trials, presented in pseudorandom order, were divided into three blocks of 240 stimuli, equally split by type. A brief training session familiarised participants with the task. Reaction times were recorded using a pedal board connected to the TriggerStation™ (BRAINTRENDS LTD 2010, Rome, Italy). Breaks were allowed between blocks to prevent fatigue and maintain focus.

### Redundant signal effect analysis

The redundant signal effect (RSE [55,56]) occurs when individuals are asked to make fast responses in a simple detection task. This effect reflects the relative gain in RTs, observed when stimuli are presented simultaneously in multiple sensory modalities (i.e., redundant stimuli), as opposed to a single modality [55–58]. We adjusted reaction time data as follows. First, responses faster than 120 msec were classified as “fast guesses” and were removed from the analysis [59]. Next, for each stimulus type and participant, we trimmed all RTs falling outside 2 SD from the mean (5.09% of data was rejected from raw RTs). Finally, we assessed the normality of data distributions using Lilliefors tests for each stimulus type, implemented in MATLAB (R2023a, MathWorks Inc., Natick, MA, USA). To quantify the redundant signal effect, we computed Friedman tests for each of the modality triplets (A/T/AT, A/V/AV, V/T/VT) using the MATLAB *friedman* function. Where necessary, post-hoc analyses were conducted by comparing each bimodal condition with its respective unimodal condition and applying Tukey-Kramer multiple comparisons correction. These comparisons were conducted using MATLAB’s *multcompare* function.

### Race model inequality analysis

Several models have been proposed to explain RSE, including the so-called *race models*. In race models, the two components of a bimodal stimulation are assumed to be processed in separate sensory channels, and the faster channel determines processing time [60,61]. Therefore, these models imply that RSE is a mere consequence of statistical facilitation or probability summation [62], overlooking superadditive enhancements provided by multisensory integration [63]. To distinguish between separated processing (race model) and integrated processing (multisensory integration), Miller (1982) derived *race model inequality* (RMI), which has become an important testing tool for the analysis of redundancy gains [60,64,65]. RMI states that the cumulative RT distribution for the redundant stimuli never exceeds the sum of RTs distribution for the unimodal stimuli, while rejection or violation of the inequality reflects multisensory integration [60,64,66]. We tested for such RMI violations by following the procedure reported in Saltafossi et al. (2023), based on Mahoney and Verghese (2019). We organised raw RTs into 21 progressively increasing time bins. This involved determining a specific RT range for each participant by subtracting their slowest RT from the fastest RT. Subsequently, we incrementally added the 5% of this range to each time bin. The cumulative distribution frequency (CDF) was then constructed by summing the total probabilities across the quantized bins, resulting in 21 time bins (0%, 0% + 5%, 0% + 5% + 10%, etc.) for each of the three multisensory pairs (AT, AV, VT). We computed the independent version of the race model [67] using the following formula:

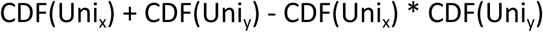

Individual RMI values were generated for each time bin by subtracting the predicted CDF (i.e., independent race model) from the actual CDF representing multisensory conditions. Violations of RMI occur when probability difference values (i.e., actual CDF - predicted CDF) are positive. However, to probe whether there was a statistically significant violation while controlling for Type I error [68], a series of permutation tests (n = 10001) were run over the violated portion (i.e., positive values) of the group-averaged difference wave, using MATLAB *rmiperm* function [66]. Results provide a Tmax value, 95% criterion Tcritic value, and p-value to determine whether multisensory integration took place across the study sample [69,70].

### Statistical analysis relating respiration and reaction times

Respiratory data were processed using custom MATLAB scripts. Respiration phase angles were extracted according to a well-validated procedure [11,18,19]. Specifically, points of peak inspiration (peaks) and expiration (troughs) were identified within the normalised (z-scored) respiratory signal using MATLAB *findpeaks* function with parameter adjustments. Then, respiration cycles were centred around peak inspiration (phase 0) through double interpolation: phase angles were linearly interpolated from trough to peak (-π to 0) and vice versa, from peak to trough (0 to π).

To relate respiration to behavioural data, we carried out bin-wise analyses on RTs collapsed across conditions (regardless of the stimulus type) and on condition-dependent RTs (i.e., unimodal and bimodal). Thus, we first partitioned the respiratory cycle into 60 equidistant and overlapping phase bins. Moving along the cycle in increments of Δω = π/30, we aggregated trimmed RTs according to the stimulus onset phase computed at a respiration angle of ω ±π/10 [11]. Respiration phase (bin)-dependent averaged RTs were obtained for each participant. Finally, RTs were z-scored and averaged across participants, resulting in the grand average phase-dependent RTs at the population level. Likewise, for each condition (unimodal and bimodal) we derived the grand average RTs.

We employed a linear mixed effect model (LMEM) to investigate whether respiration affects speeded responses, through MATLAB *fitlme* function. A first (base) model predicted RTs as a combination of the fixed effect of stimulus type (i.e., unimodal and bimodal) and the random intercept for participants. We used the following formula defined according to Wilkinson notation:

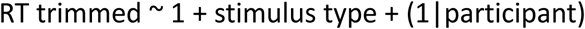

An alternative LMEM modelled the same variables adding the fixed effect of respiration angle (with separate sine and cosine contributions) as follows:

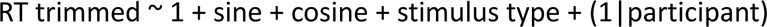

The MATLAB *compare* function was employed to contrast the two LMEMs, thereby assessing the modulatory effect of respiration on RTs through a theoretical likelihood ratio test (LRT). To further evaluate the significance of this respiration modulatory effect, the LMEM beta weights for the sine and cosine factors were combined into a respiratory phase vector norm, defined as:

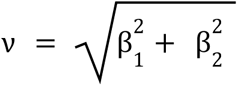

This empirical combination of sine and cosine components was tested against a null distribution generated from randomised individual RTs. Specifically, the LMEM was recalculated 1000 times, shuffling each participant’s RTs across the 60 respiratory phase bins. For each iteration, the resulting beta weights for sine and cosine were combined as described above, producing a distribution of “null vector norms”. The significance of the observed empirical vector norm was determined by computing its percentile rank within the null distribution’s density function.

To corroborate the LMEM findings and examine the directionality of the effect, the distributions of both overall RTs and condition-dependent RTs across the respiratory cycle were tested for uniformity using the Rayleigh test provided by the *circstat* toolbox [71] running on MATLAB. Only for the condition-dependent RTs, a Watson-Williams (WW) test was performed, contrasting their circular means to check whether reaction times following unimodal and bimodal trials were differentially modulated by respiration. To test for significant respiration-related changes in RTs (e.g., lowering and increasing), z-scored RTs were first transformed into t-values and then subjected to two-tailed t-tests to determine whether the observed changes in the data are unlikely to occur by random chance. Significance levels were adjusted for false discovery rate (FDR) using the MATLAB *mafdr* function.

### Statistical analysis relating respiration and multisensory integration

To investigate whether respiration played a role in shaping multisensory integration, RMI analyses were performed on 4 respiration phases, including both transition and non-transition phases. Transition phases, i.e., expiration-to-inspiration (ex2in) and inspiration-to-expiration (in2ex), were defined as ¾π to −¾π and −π/4 to π/4, respectively, while non-transition phases, i.e., ongoing inspiration and expiration, were defined as −¾π to −π/4 and π/4 to ¾π [19]. For bimodal conditions where the rejection of the inequality was observed, trials were clustered into the 4 phases according to their stimulus onset phase. As RMI analysis requires a comparable number of observations between conditions with a minimum of 20 [68], trials were down-sampled for each participant using the MATLAB *sampleDown* function provided by the *RSE-box* toolbox [72]. Additionally, participants with low number of trials (< 20) for each stimulus type and respiratory phase were excluded from the analysis (6 participants for AT-ex2in, 6 for AT-inspiration, 4 for AT-expiration, 2 for AT-in2ex, 6 for AV-ex2in, 5 for AV-inspiration, 3 for AV-in2ex, and 4 for AV-expiration). In this approach, the down-sampled raw reaction times were subjected to RMI analyses, as previously described. This process produced group-averaged RMI waveforms for each of the four phases across all bimodal conditions.

Aiming to quantify the difference in multisensory integration between respiratory phases, we calculated individual area-under-the-curve (AUC) values of RMI wave (actual CDF - predicted CDF). AUC served to determine the magnitude of multisensory integration (MMSI) as described in [54,69,73]. Therefore, RMI values from the first time bin (0%) were summed with RMI values obtained from the second time bin (5%) and divided by two (AUC 1 = (0% + 5%)/2). This process was repeated for the subsequent time bins until the last violated (positive) time bin was reached. Finally, to analyse the effect of the respiratory phase on the magnitude of multisensory integration, we set up a model comparison of two LMEMs for each bimodal condition. The first (base) model, including only the effect of the AUC window and the random effect of participants, was defined (in Wilkinson notation) as:

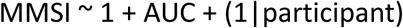

We entered as many AUCs as long as at least one phase still had a positive RMI value (from AUC 1 to AUC 11). The second model added to the previous one the categorical information of the respiratory phase (ex2in, inspiration, in2ex, expiration):

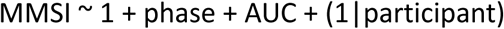

To substantiate whether the addition of the respiratory information improved the model fit, we contrasted LMEMs with and without the respiratory phase predictor using theoretical LRT provided by the MATLAB *compare* function. ANOVAs were performed to determine if the coefficient estimates were significantly different from zero. Last, KW tests and multiple comparisons corrected post-hocs were run, using MATLAB *kruskalwallis* and *multcompare* (with Tukey-Kramer critical value type) functions, respectively, to show which phase was associated with greater MMSI.

### Statistical analysis of the alignment between respiration and response onsets

To investigate whether participants temporally adjusted their respiration rhythm to response onsets, we first extracted respiration phases of the response onsets for all trials (regardless of the stimulus type) and then separately for unimodal and bimodal trials. Circular statistics (Rayleigh tests, *circstat* toolbox [71]) were applied to test the hypothesis that, across participants, response onsets-averaged respiratory phase values were not distributed uniformly across the cycle.

## Results

### Redundant signal effect

We recorded RTs during a simple detection task in which supra-threshold unimodal (A, T, V) and bimodal (AT, AV, VT) stimuli were presented. Across participants, we observed an average hit rate above the 85% for all stimulus types (see Supplementary, Table 1), which was in line with previous literature positing ceiling performance as an assumption for the RMI analysis [72].

Since each stimulus type’s data (RTs) did not follow the normal distribution (Lilliefors tests; A: *p* = .001; T: *p* = .002; V: *p* = .005; AT: *p* = .006; AV: *p* = .005; VT: *p* = .002) (further statistics are reported in the Supplementary, Table 2), we investigated whether RSE was observed in each cross-modal combination using Friedman and Tukey-Kramer post-hoc tests (full results tables are reported in the Supplementary, Tables 3 - 5). For A/T/AT F-test, stimulus type affected RTs (χ²(2, 78) = 72.80, *p* < .001, W = 0.91), and responses to AT stimuli were faster than those to unimodal stimuli (vs. A, *p* < .001; vs. T, *p* < .001). Also, unimodal A RTs were significantly faster compared to T RTs (p < .001) (**Figure 2a**). Likewise, A/V/AV F-test revealed a significant association of stimulus type and RTs (χ²(2, 78) = 60.80, *p* < .001, W = 0.76). Tukey-Kramer post-hoc confirmed RSE, with AV RTs being faster than both A and V RTs (vs. A, *p* < .001; vs. V, *p* < .001) (**Figure 2b**). Last, for V/T/VT F-test, the effect of stimulus type was significant (χ²(2, 78) = 61.85, *p* < .001, W = 0.73), with Tukey-Kramer post-hoc showing faster RTs to bimodal compared to unimodal stimuli (vs. T, *p* < .001; vs. V, *p* = .003). Moreover, responses to V stimuli were faster than responses to T stimuli (*p* < .001) (**Figure 2c**). In summary, bimodal stimulations lead to faster responses compared to unimodal stimulations. This finding prompted further analysis to investigate whether the observed response speed enhancement is attributable to multisensory integration.

**Figure 2.**
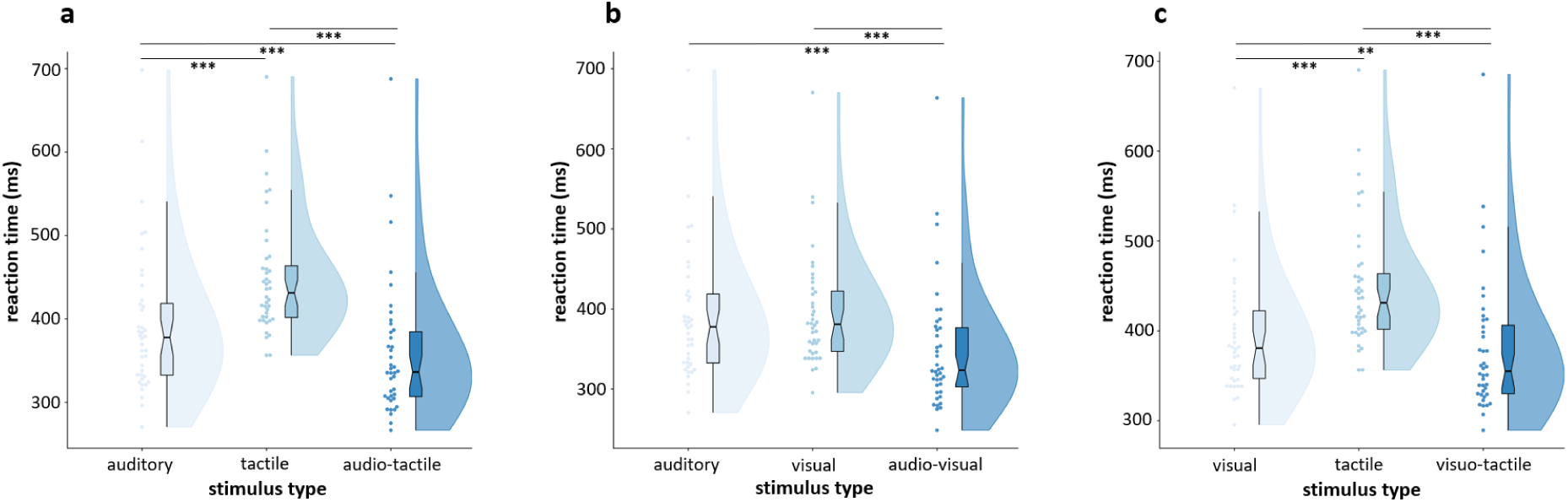
Redundant signal effect. Raincloud plots showing the behavioural facilitation related to bimodal stimulations (RSE) for each modality triplets A/T/AT (**a**), A/V/AV (**b**), and V/T/VT (**c**). Lines with three and two asterisks represent *p* < .001 and *p* < .01, respectively.

### Race model inequality and multisensory integration

To test for the presence of multisensory integration, rather than only RSE, we employed Gondan’s permutation tests [73], which iteratively compare CDFs from actual data with CDFs from the race model. AT RMI was violated within time bins ranging from the 2^nd^ to the 8^th^ (*Tmax* = 4.92, *Tcritic* = 2.20, *p* < .001) (**Figure 3a**). Similarly, AV RMI was violated over time bins 2^nd^ to 7^th^ (*Tmax* = 6.42, *Tcritic* = 2.08, *p* < .001) (**Figure 3b**). In contrast, VT didn’t show multisensory integration, as permutation tests over the 3^rd^ and 4^th^ time bins returned rejection of the alternative hypothesis (*p* > .05) (**Figure 3c**). While these results are confirmatory rather than novel, they pose a prerequisite for investigating respiration-related changes in multisensory integration.

**Figure 3.**
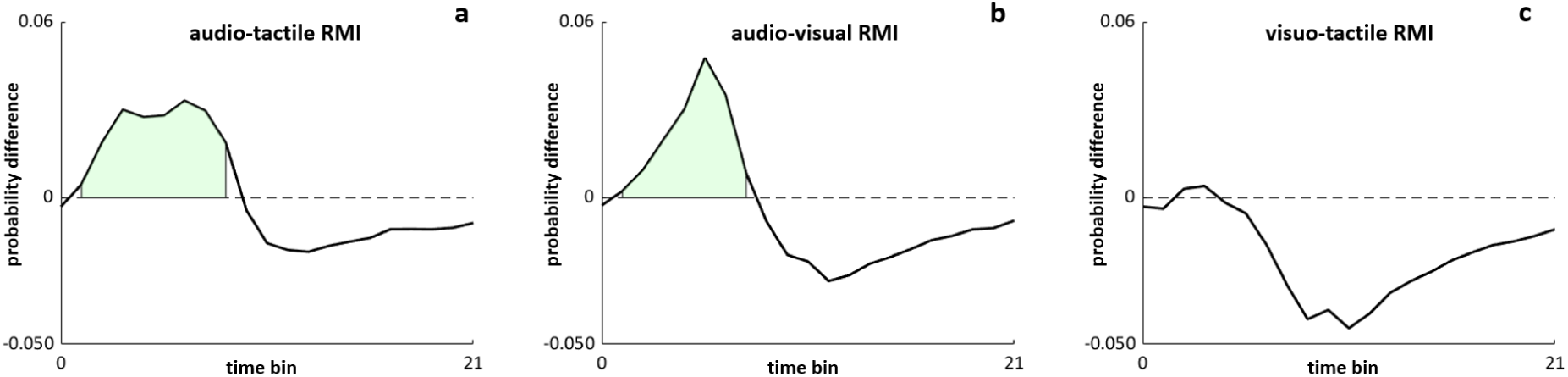
Race model inequality and multisensory integration. Group-averaged probability CDFs difference (actual – predicted) across all time bins (from 0 to 21) are depicted for each bimodal condition. Light green areas represent the violated portion of the waveform (RMI), indicative of multisensory integration (**a**, **b**).

### Respiratory modulation of reaction times

We assessed whether participants’ performance systematically varied over the respiratory cycle. We first demonstrated that incorporating respiratory phase sine and cosine terms into the full model, along with stimulus type information, significantly improved the fit compared to the base model (LRT-stat χ²(2) = 133.16, *p* < .001) (all models stats are reported in the Supplementary, Tables 6 - 8). Furthermore, the empirical vector norm, derived from combining the LMEM beta weights for sine and cosine, exceeded all null vector norms derived from 1000 iterations of randomised RTs across the 60 bins (*p* < .001) (see Supplementary, Figure S1).

Then, two separated respiratory phase bin-wise analyses resulted in RT group distributions across the respiratory cycle, for trials pooled together (regardless of the stimulus type) and for trials separated based on the stimulus type (unimodal vs bimodal). Through circular statistics, we confirmed the hypothesis that pooled RTs varied with respiration, regardless of the stimulus type. This was substantiated by the non-uniform distribution of RTs (*Z*_Rayleigh_ = 4.63, *p* = .009), marked by a general increase towards the ex2in phase (*V*_mean_ = 3.02, 95% CI [3.70 2.34]) (**Figure 4a**). Importantly, respiration-related changes in RTs were consistent across our sample: 32 participants out of 40 presented this effect (*p* < .05) (the full list is reported in the Supplementary, Table 9). Examining the time-course of this modulation revealed slower responses within the ex2in transition (corresponding to bins’ angles: −3.14 to −2.72, 2.29 to 3.14 rad, *p*_FDR_ < .05), while faster responses were found roughly around peak inspiration and during early expiration (corresponding to bins’ angles: −2.18 to −1.44, −1.33, and 0.05 to 1.44 rad respectively, *p*_FDR_ < .05) (**Figure 4a**) (see Supplementary for t-values and relative FDR adjusted p-values, Table 10).

**Figure 4.**
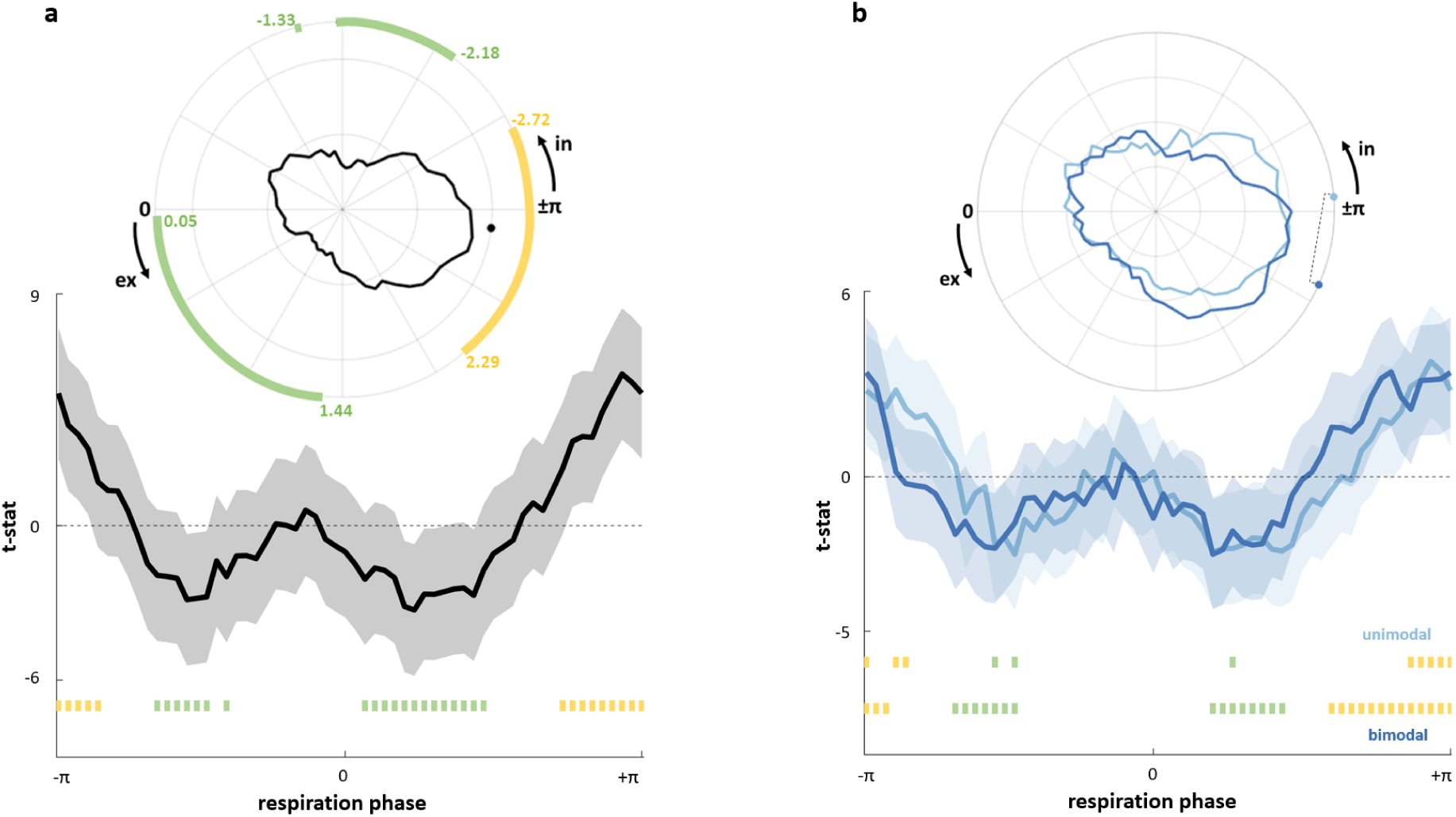
Respiratory modulation of reaction times. (**a**) Polar visualisation shows overall RTs across the respiration cycle. A Solid black line illustrates respiration phase-dependent RTs, black dot indicates the circular mean. Slower RTs occur in the ex2in phase (yellow markings *p*_FDR_ < .05), while faster RTs align with peak inspiration and early expiration (green markings *p*_FDR_ < .05). The 2D line graph displays t-stat values across respiration, with shaded areas for standard deviation and green/yellow squares for significant RT changes. (**b**) RTs-respiration phase comparison for unimodal (light blue) and bimodal trials (blue). In the polar plot, color-matched dots mark circular means, connected by a dashed line for a significant Watson-Williams test (p < .05). The 2D graph shows t-stat values, with color-coded shaded areas for standard deviation. Green and yellow squares indicate significant RT changes for each stimulation type.

In the second circular bin-wise analysis, both responses to unimodal and bimodal stimuli exhibited non-uniform distributions across the respiratory cycle (unimodal *Z*_Rayleigh_ = 3.26, *p* = .036; bimodal *Z*_Rayleigh_ = 3.15, *p* = .040), but the circular means of unimodal (*V*_mean_ = −3.06, 95% CI [−2.20 −3.92]) and bimodal stimuli (*V*_mean_ = 2.72, 95% CI [3.60 1.84]) differed significantly from each other (WW = 10.03, *p* = .002) (**Figure 4b**). A significant increase of RTs took place within the ex2in transition for both unimodal (bins’ angles: −3.14, −2.82 to −2.72, 2.72 to 3.14 rad, *p*_FDR_ < .05) and bimodal trials (bins’ angles: −3.14 to −2.93, 1.86 to 3.14 rad, *p*_FDR_ < .05), while lower RTs were mainly located towards peak inspiration and early expiration, again for both unimodal (bins’ angles: −1.76, −1.54, 0.80 rad, *p*_FDR_ < .05) and bimodal trials (bins’ angles: −2.18 to −1.54, 0.59 to 1.33 rad, *p*_FDR_ < .05) (**Figure 4b**) (see Supplementary for t-values and relative FDR adjusted p-values, Table 10). Respiratory modulations of RTs were observed in 34 participants out of 40, for both unimodal and bimodal stimuli (*p* < .05) (the full list is reported in the Supplementary, Table 9).

### Respiratory modulation of multisensory integration

Since the previous analysis had established a meaningful modulation of reaction times as a function of the respiration phase, we next addressed the question of whether respiratory phases would also modulate the magnitude of multisensory integration. To address potential changes in multisensory integration due to respiration, we set up two LMEMs for the bimodal conditions that demonstrated multisensory integration, specifically AT and AV stimuli, as detailed above. The base model, featuring only the AUC predictor, was contrasted with a second LMEM, which included the respiratory phase information.

Regarding AT, the model comparison between the two LMEMs confirmed that including phase information significantly improved the model fit (LR-stat χ²(3) = 38.27, *p* < .001) (see Supplementary for the models’ stats, Tables 11 - 13). The model yielded significant effects of both the AUC (in a time window reaching AUC 11) (*F*(10, 1548) = 4.11, *p* < .001) and the respiratory phase (*F*(3, 1548) = 12.92, *p* < .001). Respiratory phase clearly influenced AT integration (KW χ²(3) = 35.13, *p* < .001), with Tukey-Kramer post-hoc revealing that ex2in (vs. inspiration mean difference = 137.89, *p* < .001; vs. in2ex mean difference = 161.83, *p* < .001) and expiration (vs. inspiration mean difference = 101.39, *p* = .01; vs. in2ex mean difference = 125.32, *p* < .001) exerted the greatest impact (**Figure 5a**).

**Figure 5.**
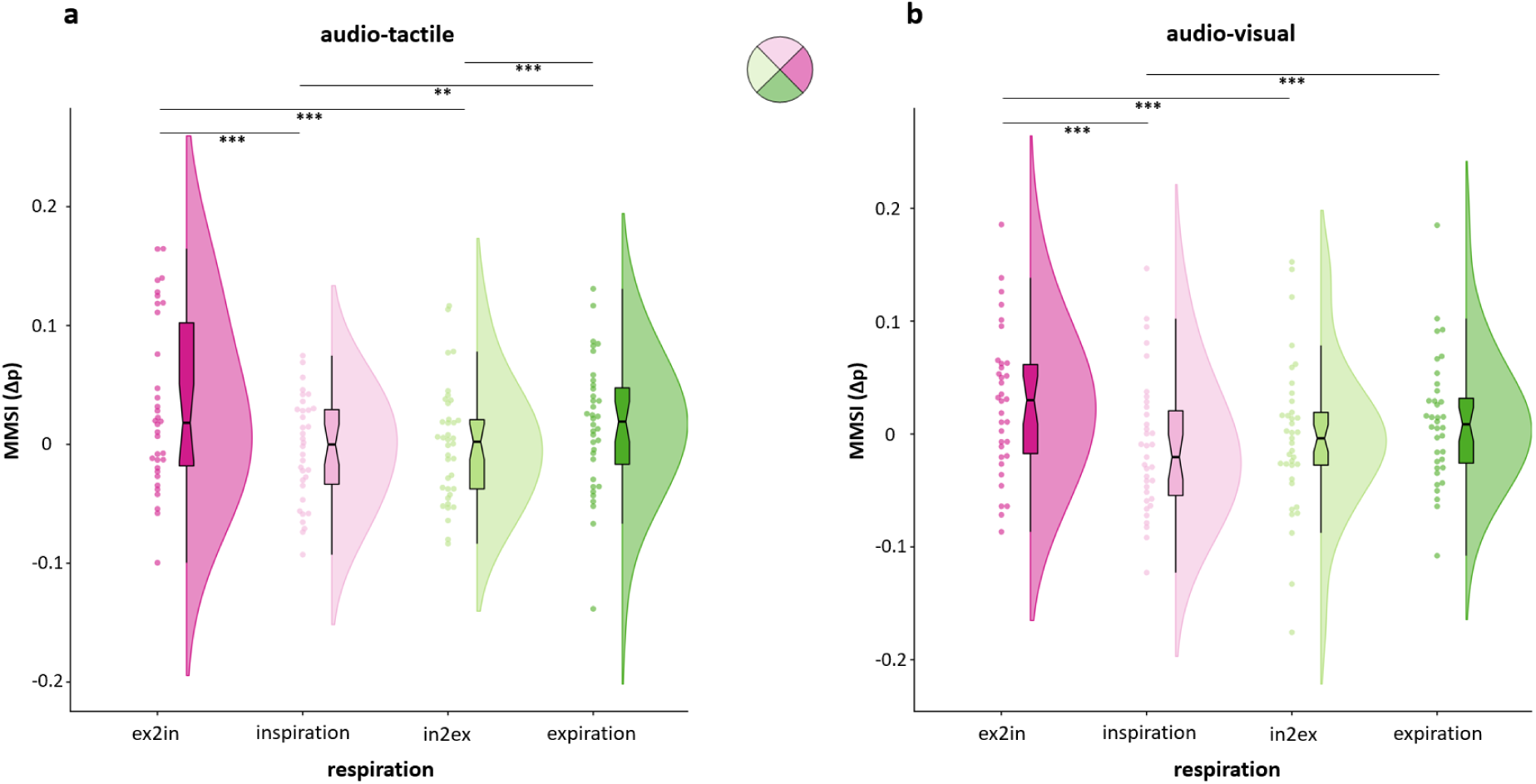
Respiratory modulation of multisensory integration. Both AT (**a**) and AV (**b**) magnitude of multisensory integration particularly increased in ex2in and expiration phases, as illustrated by phase-matched coloured raincloud plots. Black lines and asterisks represent significant post-hoc comparisons: ** *p* = .01, *** *p* < .001.

In a similar vein, for the AV stimulations, the model including the respiratory phases provided the best fit (LR-stat χ²(3) = 36.50, *p* < .001) (see Supplementary for the models’ stats, Tables 14 - 16). While there was no effect of AUC (up to AUC 10), the respiratory phase significantly modulated multisensory integration as constantly shown by both ANOVA (*F*(3, 1407) = 12.32, *p* < .001) and KW test (KW χ²(3) = 44.21, *p* < .001). Tukey-Kramer post-hoc confirmed that this effect was mostly driven by ex2in (vs. inspiration mean difference = 195.26, *p* < .001; vs. in2ex mean difference = 138.21, *p* < .001) and, to a lesser extent, by expiration (vs. inspiration mean difference = 125.13, *p* < .001) (**Figure 5b**). Overall, these findings indicate a clear role of respiration in shaping multisensory integration, as adding the phase information to the models greatly improved their explanatory power. Moreover, specific respiratory phases seem to boost multisensory integration, suggesting potential interactions with the neural processes underpinning this perceptual feature.

### Respiratory cycle is aligned with response onset

A growing number of empirical findings point to a role of respiration in shaping information sampling and behaviour (for recent reviews, see [9,24,26]). Therefore, we directly tested the association between the respiratory phase and response onsets, as participants may potentially adjust their breathing patterns to meaningful paradigm events. Results showed that, overall, the average phase angles were highly clustered towards expiration and deviated significantly from the null hypothesis of a uniform distribution (*Z*_Rayleigh_ = 17.78, *p* < .001, *V*_mean_ = 0.60, 95% CI [0.90 0.30]) (**Figure 6a**). Inspecting individual data revealed that 24 participants out of 40 systematically aligned their respiratory behaviour around response time (the full list is reported in the Supplementary, Table 17). We tested for such an alignment also by splitting data according to the stimulation type (unimodal and bimodal). Circular statistics returned a significant grouping of response onsets average phase angles again within the expiration phase for both unimodal (*Z*_Rayleigh_ = 14.40, *p* < .001, *V*_mean_ = 0.64, 95% CI [0.99 0.30]) and bimodal trials (*Z*_Rayleigh_ = 14.14, *p* < .001, *V*_mean_ = 0.53, 95% CI [0.88 0.19]) (**Figure 6b**). These effects were consistent across our study sample, as 17 and 18 participants (out of 40) adapted their respiration to response onset for unimodal and bimodal trials, respectively (*p* < .05) (the full list is reported in the Supplementary, Table 17). Taken together, this analysis provides additional evidence of the intricate interplay between respiration and perception. While RTs were found to be higher within the ex2in and lower during inspiration and early expiration, as well as response onsets clustering towards expiration, the ex2in transition would have benefited multisensory integration in both AT and AV conditions.

**Figure 6.**
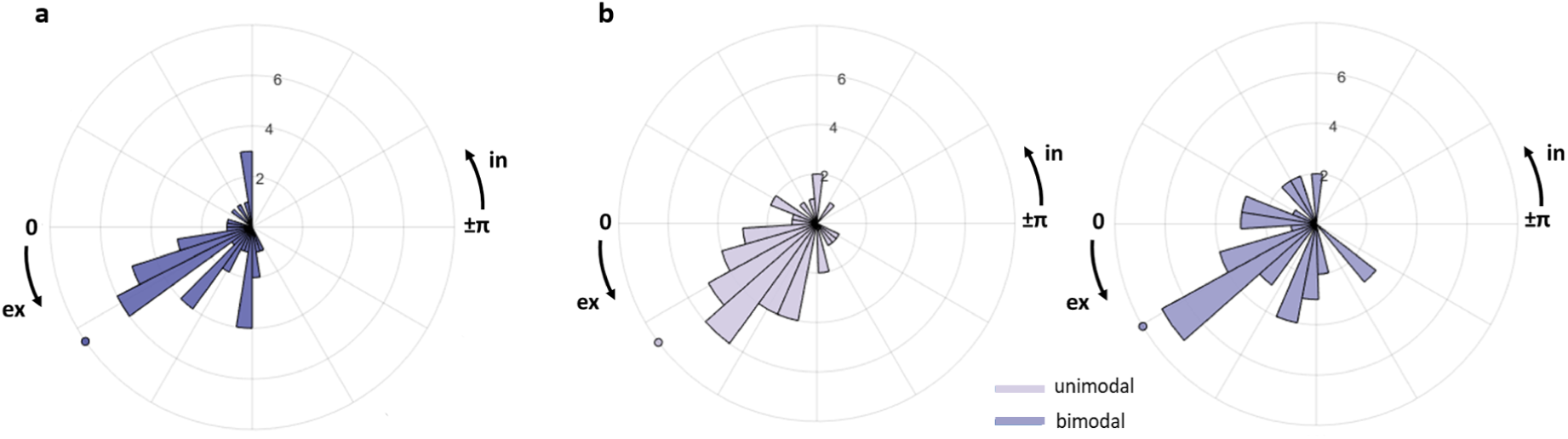
Respiratory cycle is aligned with response onset. (**a**) Distribution of participants’ average phase values at response onset, pooled across stimulus types, shows a preference for early expiration (coloured dot: circular mean). (**b**) Polar histograms reveal response onset clustering toward early expiration for both unimodal and bimodal stimulation (colour-matched dots: circular means).

## Discussion

To address the relationships between respiration and multisensory integration, we tested whether and how respiratory phases relate with various facets of multisensory perception, such as RTs, MMSI, and response onsets to unimodal and bimodal stimuli. We were able to demonstrate that respiration modulates speeded responses and multisensory integration, with participants timing their responses to align with their breathing rhythm.

### Distinct time windows within the respiratory cycle speed up reaction times

While research on cardiac interoception has long highlighted the distinct roles of systole and diastole in regulating various cognitive functions [74], including sensory [54,75–81] and motor processes [82–86], research on respiration is limited. Here, by employing LMEMs and circular statistics, we showed a complex picture of the interplay between the breathing cycle and perception among multiple sensory modalities. Notably, responses were speeded up within two time-windows of the cycle, namely around peak inspiration and early expiration. Despite earlier inconclusive outcomes [31,87], recent investigations on visual information processing have demonstrated that best performances are achieved during inspiration, with some offering neural evidence based on olfactory bulb-mediated phase-amplitude coupling to support these effects [11,27,29,30]. Various findings have been reported regarding the auditory system. For instance, two studies indicated that the effects of respiration emerged only when participants were required to exert some form of control over their breathing.

One study [32] found this resulted in prolonged RTs, while another [35] observed faster startle responses to auditory stimuli during expiration. In contrast, somatosensory perception appears to benefit most from later phases of the respiratory cycle. Specifically, the accurate detection of near-threshold tactile stimuli was enhanced immediately after the onset of expiration [36]. Despite the variability introduced by differing methods for monitoring respiration (e.g., belts, manometers, or temperature sensors), the specific breathing instructions provided to participants (spontaneous versus controlled), the physical characteristics of the stimuli (near-threshold versus suprathreshold), and the diverse approaches used to assess respiration-behavior relationships, our findings align with previous research. Together, they highlight distinct time windows, within the respiratory cycle, where exteroceptive processing is enhanced.

### The expiration-to-inspiration transition slows down reaction times

The other main finding of our study is that speeded responses to both types of stimuli (i.e., unimodal and bimodal) were slowed down during the transition from expiration to inspiration (ex2in). Similarly, a decrease in memory-related behavioural performance has been pointed out during respiratory phase transitions [88,89]. Recall accuracy and RTs to test cues were significantly worsened in a delayed matching-to-sample task when the transition from expiration to inspiration occurred during the retrieval process [88]. A neuroimaging study by Nakamura et al. (2022) demonstrated that this ex2in transition during retrieval negatively modulates metabolic responses in critical hubs of the ventral attention network [90,91], including the anterior right temporoparietal junction, right middle frontal gyrus, and dorsomedial prefrontal cortex. In addition to the olfactory bulb-mediated pathways mentioned earlier, other mechanisms have been proposed to explain how feedback from pontomedullary respiratory centers influences large-scale brain dynamics during the ex2in phase [92]. For instance, GABAergic inhibition of glutamatergic neurons in the parabrachial nucleus during ex2in downregulates activity in the basal forebrain, hippocampus, and upstream cortical regions [93,94]. Moreover, the rhythmic firing of the PreBötzinger Complex (PreBötC), the primary generator of inspiration [7,95], modulates hippocampal ensemble dynamics and memory performance via abrupt signals evoked during ex2in transitions [96,97]. While previous research has focused on attentional and memory processing, we hypothesise that similar neural pathways underlie the ex2in-locked reductions in RTs observed in our study. Interestingly, both unimodal and bimodal stimuli followed similar respiratory modulation patterns, yet their circular means differed significantly. Functionally, both stimuli were framed within the ex2in phase, although the extent to which respiration uniquely impacts these two sensory processing types remains uncertain.

### Multisensory integration is enhanced during the expiration-to-inspiration transition

Information coming from multiple senses is combined in our brains to yield multimodally determined percepts [98]. This sensory richness provides behavioural advantages, as demonstrated in our study by the Redundancy Signal Effect (RSE). All three crossmodal combinations resulted in faster RTs compared to those observed with single-modality stimuli. However, when examining the potential super-additive interactions driving these redundancy gains, only the AT and AV combinations showed evidence of multisensory integration. In contrast, the RMI analysis did not reveal clear evidence of VT integration across different time windows, possibly due to a ceiling effect limiting VT interactions. Multisensory integration typically arises when individual sensory stimuli are weak in eliciting a response on their own, a phenomenon known as ‘inverse effectiveness’ [99–101]. Here, the high effectiveness or low variability of V and T stimuli when combined may have diminished the likelihood of VT integration.

LMEMs were applied to multisensory integration (MMSI) indices calculated for the AT and AV combinations. Once again, respiration - categorized into phase transitions (in2ex and ex2in) and non-transition phases (inspiration and expiration) - was found to predict the MMSI indices, our behavioural proxy for multisensory integration. Unlike RTs facilitation, AT and AV multisensory integration were predominantly enhanced during the ex2in phase, while expiration had a lesser or secondary influence compared to other phases. These findings suggest that respiration plays a role as a physiological modulator of brain activity, potentially influencing neural processes associated with multisensory integration. Specifically, it may act as a ‘rescue’ mechanism during slower responses to bimodal stimuli [102]. Relatedly, it has been recently demonstrated that both oscillatory and non-oscillatory brain activity covary with the respiratory cycle. For instance, non-oscillatory fluctuations in cortical excitability, driven by respiratory phases, are marked by changes in the excitation-inhibition (E-I) balance, with steepest slopes, indicative of inhibitory dominance (E<I), around the ex2in transition [18]. Multisensory binding also relates to temporal and scale-free brain activity dynamics [103] and to the balance of glutamate and GABA concentrations, which, along with genetic variation in related genes, influence individual variability in AT multisensory integration [104]. In auditory-visual paradigms, for example, GABA concentration in the superior temporal sulcus enhances gamma-band power and AV perceptual binding [105]. We propose that the ex2in phase fine-tunes brain activity to align with optimal excitability states (E<I), facilitating multisensory integration.

Our findings speak to recent theoretical proposals like body-extended multisensory integration, scaffolding, and predictive processing [106], as well as to the multi-timescale conceptualisation of multisensory integration [107]. In this context, slow rhythms like respiration act as carrier waves, regulating higher cortical dynamics while establishing temporal precision [9,10,108,109]. As a consequence, prestimulus functional coupling across both frequency bands and brain regions [110–113], and fluctuations in ongoing oscillations that support multisensory integration [114–116] may be intrinsically orchestrated by respiration. This coordination might impose specific brain states within which multisensory processing occurs.

### Response onsets cluster within early expiration

Building on concepts like embodied predictive interoception coding (EPIC) [117] and interoceptive active inference [10,118] which constraint model updating and PEs minimisation to visceral ascending and descending information, our results suggest that response timing is intricately linked to specific respiratory phases, particularly during early expiration. Given respiration’s predictability and adaptability [119], our findings highlight how sensory sampling and environmental interactions align with specific brain-body states [11,120–122]. This matches with the active sensing hypothesis, and experimental results showing that task, stimulus, response onsets, and actions triggered externally or through mental imagery synchronise with respiration [23,27,29,36,39,40,46,123].

While we cannot definitively establish directionality between response clustering and RTs reduction, these phenomena likely reflect respiration-based precision-weighting mechanisms. In predictive processing, precision reflects confidence in predictions and prediction errors, modulated via neurotransmitter activity [124–127]. Notably, brainstem breathing circuits directly elicit a noradrenergic release from the locus coeruleus, tightly coupling respiration to arousal and behavioural regulation [128–132]. This may explain the cognitive and attentional benefits observed with controlled breathing practices in traditions like yoga and martial arts, where respiratory regulation enhances focus and clarity [25,133–135].

## Conclusion

In conclusion, our findings strongly suggest that respiration actively shapes exteroceptive processing, encompassing both unisensory and multisensory inputs. Future research should aim to disentangle the respiratory entrainment of responses from genuine multisensory integration, while also identifying the specific neurophysiological mechanisms supporting both processes. This would help substantiate a more comprehensive perspective on multisensory integration extending to an embodied hierarchical neuroarchitecture in which slower bodily rhythms play a crucial role [3,136–139]. Such insights could also advance our understanding of neurological and psychiatric disorders characterised by disrupted interoceptive processing [19,140–144].

## Acknowledgments

We would like to thank Eleonora Pozzi and Serena Turchi for their assistance during data collection. We are also grateful to Başak Bayram for her insightful discussions that contributed to the development of this paper. D.S.K. is supported by the IMF (KL 1 2 22 01) and the DFG (KL 3580/1 - 1). M.C. is supported by Boosting Ingenium for Excellence (BI4E) funded by the European Union’s HORIZON-WIDERA-2021-ACCESS-05-01-European Excellence Initiative under the Grant Agreement No. 101071321, and PNRR Project “Boost for Interdisciplinarity” (“NextGenerationEU”, “MUR-Fondo Promozione e Sviluppo - DM 737/2021”, INTRIGUE). F.F. is supported by: PRIN 2022 - Interoception and Active Aging (InterActing) - Prot. 2022JS4SY2, PRIN 2022 PNRR Project “Metaphor and epistemic injustice in mental illness: the case of schizophrenia” - CUP D53D23020890001, and “Departments of Excellence 2023-2027” initiative of the Italian Ministry of University and Research for the Department of Neuroscience, Imaging and Clinical Sciences (DNISC) of the University of Chieti-Pescara. We acknowledge support from the Open Access Publication Fund of the University of Münster.

## Author contributions

Conceptualisation F.F., M.C., A.Z.; Methodology A.Z., D.S.K., M.G.P., M.S.; Investigation M.S.; Writing – original draft M.S.; Writing – review & editing M.S., A.Z., D.S.K., M.G.P., F.F., M.C.; Visualisation M.S. D.S.K.; Funding acquisition F.F., M.C..

## Data availability

The data that support the findings of this study are available from the corresponding author upon reasonable request.

## Code availability

The code used for statistical analyses and result visualisations in this study is available from the corresponding author upon request. The analyses were performed using MATLAB R2023a and RStudio 2023.12.0.

