## Supplementary for "Respiration facilitates behaviour during multisensory integration"

### 1. Experimental setup - stimuli and thresholding procedure description

The stimulus A was a pure tone (1000 Hz; 30 msec of duration) presented approximately at 60 dB using a buzzer; V stimulus (30 msec of duration) was delivered through a light-emitting diode (LED) (5 mm diameter; 200 mcd); T stimulus consisted of a suprathreshold electrical pulse (100 µs of duration), delivered on the middle finger of the right hand using a Digitimer

(DS7A, Digitimer Ltd., Welwyn Garden City, UK). A and V stimuli were presented through an in-house box containing a fixation cross, the buzzer, and the LED (main text: **Figure 1a**). Stimuli presentation was controlled with the E-Prime 3.0 software (Psychology Software Tools, Pittsburgh, PA, USA) connected to a TriggerStation™ (BRAINTRENDS LTD 2010, Rome, Italy). Individual thresholds of the tactile stimulus were set to be clearly suprathreshold using the method of the limits [1]. Before starting the experiment, the intensity of the stimulator was set to 0 mA and then progressively increased by 1 mA until the subject reported to clearly perceive the stimulation. Then, the participant was additionally stimulated 5 times: as soon as one of the additional stimuli was not detected, the intensity was increased by 1 mA, and the procedure was repeated [2,3].

### Table 1. Accuracy

| **stimulus type** | **mean accuracy (%)** |
|---|---|
| A | 97.95 % |
| V | 97.14 % |
| T | 86.28 % |
| AT | 98.80 % |
| AV | 99.10 % |
| VT | 98.50 % |

### Table 2. Lilliefors tests for normality

| **stimulus type** | **decision for null** | **p-value** | **k-stat** | **critical value** |
|---|---|---|---|---|

| | hypothesis | | | |
|---|---|---|---|---|
| A | h = 1 | .001 | 0.19 | 0.14 |
| V | h = 1 | .027 | 0.15 | 0.14 |
| T | h = 1 | .002 | 0.18 | 0.14 |
| AT | h = 1 | .006 | 0.17 | 0.14 |
| AV | h = 1 | .005 | 0.17 | 0.14 |
| VT | h = 1 | .002 | 0.18 | 0.14 |

### Friedman's tests for Redundant Signals Effect

Table 3. A/T/AT F-test

| source | SS | DF | MS | Chi-sq | p-value |
|---|---|---|---|---|---|
| columns | 72.8 | 2 | 36.4 | 72.8 | < .001 |
| error | 7.2 | 78 | 0.09 | | |
| total | 80 | 119 | | | |

Table 3.a post hoc (Tukey-Kramer)

| stimulus type | stimulus type | low limit | difference | upper limit | p-value |
|---|---|---|---|---|---|
| T | A | 0.27 | 0.80 | 1.32 | < .001 |
| T | AT | 1.37 | 1.90 | 2.42 | < .001 |
| A | AT | 0.57 | 1.10 | 1.62 | < .001 |

### Table 4. A/V/AV F-test

| source | SS | DF | MS | Chi-sq | p-value |
|---|---|---|---|---|---|
| columns | 60.8 | 2 | 30.4 | 60.8 | < .001 |
| error | 19.2 | 78 | 0.25 | | |
| total | 80 | 119 | | | |

Table 4.a post hoc (Tukey-Kramer)

| stimulus type | stimulus type | low limit | difference | upper limit | p-value |
|---|---|---|---|---|---|
| V | A | -0.32 | 0.20 | 0.72 | .643 |
| V | AV | 1.07 | 1.60 | 2.12 | < .001 |
| A | A | 0.87 | 1.40 | 1.92 | < .001 |

### Table 5. V/T/VT F-test

| source | SS | DF | MS | Chi-sq | p-value |
|---|---|---|---|---|---|
| columns | 61.85 | 2 | 30.92 | 61.85 | < .001 |
| error | 18.15 | 78 | 0.24 | | |
| total | 80 | 119 | | | |

Table 5.a post hoc (Tukey-Kramer)

| stimulus type | stimulus type | low limit | difference | upper limit | p-value |
|---|---|---|---|---|---|
| T | V | 0.50 | 1.02 | 1.54 | < .001 |

| | | | | | |
|---|---|---|---|---|---|
| T | VT | 1.22 | 1.75 | 2.27 | < .001 |
| V | VT | 0.20 | 0.72 | 1.24 | < .001 |

**Linear Mixed Effect Models (LMEMs) for RTs and respiration**

First-base model = clean RTs ~ 1 + stimulus type + (1|participant)

Table 6. model fit stats

| **AIC** | **BIC** | **Log-Likelihood** | **deviance** |
|---|---|---|---|
| 38277 | 38303 | -19135 | 38269 |

Table 6.a fixed effects coefficients

| **name** | **estimate** | **SE** | **tStat** | **DF** | **p-value** | **lower** | **upper** |
|---|---|---|---|---|---|---|---|
| intercept | 459.3 | 11.82 | 38.87 | 4798 | < .001 | 436.13 | 482.46 |
| stimulus type | -48.63 | 0.36 | -133.85 | 4798 | < .001 | -49.347 | -47.92 |

Alternative-full model = clean RTs ~ 1 + stimulus type + sine + cosine + (1|participant)

Table 7. model fit stats

| **AIC** | **BIC** | **Log-Likelihood** | **deviance** |
|---|---|---|---|
| 38148 | 38187 | -19068 | 38136 |

Table 7.a fixed effects coefficients

| **name** | **estimate** | **SE** | **tStat** | **DF** | **p-value** | **lower** | **upper** |
|---|---|---|---|---|---|---|---|
| intercept | 410.61 | 11.80 | 34.78 | 4796 | < .001 | 387.47 | 433.76 |

| | | | | | | | |
|---|---|---|---|---|---|---|---|
| stimulus type | -48.63 | 0.36 | -135.74 | 4796 | < .001 | -49.34 | -47.93 |
| cosine | -2.85 | 0.25 | -11.34 | 4796 | < .001 | -3.34 | -2.36 |
| sine | .65 | 0.26 | 2.53 | 4796 | .012 | 0.14 | 1.15 |

Table 8. Model comparison: Theoretical Likelihood Ratio Test

| **model** | **DF** | **AIC** | **BIC** | **Log-Likelihood** | **LR-Stat** | **delta df** | **p-value** |
|---|---|---|---|---|---|---|---|
| base | 4 | 38277 | 38303 | -19135 | | | |
| full | 6 | 38148 | 38187 | -19068 | 133.16 | 2 | < .001 |

Figure S1. Alternative-full model empirical vs null distributions

Histogram shows the empirical LMEM beta for the respiratory vector norm against the null distribution computed from 1000 iterations of randomized RTs vectors on the subject level.

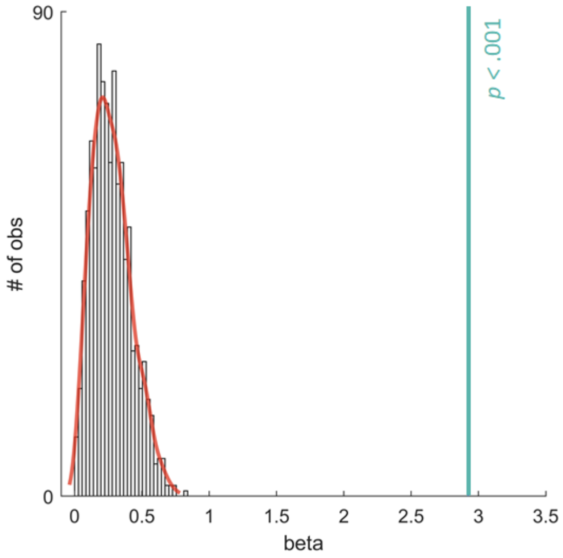

### Circular Analyses (RTs)

Table 9. Consistency across participants (RTs)

The following table lists overall, unimodal, and bimodal Z-stats from Rayleigh tests, one for each participant (from 1 to 40), with corresponding p-values. The red filling represents significant (i.e. $p < .05$) non-uniform distribution of the data.

| **Rayleigh Z** | **p-value** | **unimodal** | | **bimodal** | |
|---|---|---|---|---|---|
| 1.73 | .177 | **Rayleigh Z** | **p-value** | **Rayleigh Z** | **p-value** |
| 8.34 | < .001 | 6.17 | .002 | 3.11 | .045 |
| 11.18 | < .001 | 6.08 | .002 | 12.42 | < .001 |
| 3.33 | .036 | 12.95 | < .001 | 4.99 | .007 |
| 7.84 | < .001 | 6.82 | .001 | 5.56 | .004 |
| 6.32 | .002 | 9.73 | < .001 | 3.31 | .036 |
| 10.62 | < .001 | 5.45 | .004 | 3.4 | .033 |
| 16.57 | < .001 | 3.68 | .025 | 12.72 | < .001 |
| 13.92 | < .001 | 15 | < .001 | 4.45 | .011 |
| 2.88 | .056 | 11.94 | < .001 | 16.18 | < .001 |
| 8.98 | < .001 | 2.56 | .077 | 5.75 | .003 |
| 12.17 | < .001 | 3.48 | .031 | 3.93 | .019 |
| 6.41 | .002 | 11.53 | < .001 | 2.23 | .107 |
| 14.21 | < .001 | 5.3 | .005 | 3.14 | .043 |
| 8.53 | < .001 | 13.43 | < .001 | 5.37 | .004 |
| 2.54 | .079 | 11.41 | < .001 | 0.5 | .605 |
| 7.88 | < .001 | 4.73 | .009 | 6.42 | .002 |
| 8.92 | < .001 | 13.93 | < .001 | 3.36 | .034 |
| 15.05 | < .001 | 7.58 | < .001 | 5.45 | .004 |
| 11.26 | < .001 | 12.55 | < .001 | 3.84 | .021 |

| | | | | | |
|---|---|---|---|---|---|
| 2.28 | .102 | 13.16 | < .001 | 5.75 | .003 |
| 5.88 | .003 | 10.67 | < .001 | 3.15 | .043 |
| 4.59 | .01 | 4.66 | .009 | 15.61 | < .001 |
| 6.62 | .001 | 2.35 | .095 | 11.35 | < .001 |
| 4.45 | .012 | 1.23 | .292 | 11.62 | < .001 |
| 7.13 | .001 | 5.76 | .003 | 5.97 | .002 |
| 5.71 | .003 | 2.85 | .058 | 8.88 | < .001 |
| 9.77 | < .001 | 9.57 | < .001 | 1.26 | .284 |
| 0.3 | .743 | 10.45 | < .001 | 9.52 | < .001 |
| 2.52 | .08 | 9.95 | < .001 | 7.24 | .001 |
| 5.69 | .003 | 3.84 | .021 | 0.5 | .606 |
| 19.19 | < .001 | 4.32 | .013 | 5.73 | .003 |
| 3.92 | .02 | 17.22 | < .001 | 13.99 | < .001 |
| 10.13 | < .001 | 6.44 | .002 | 0.11 | .896 |
| 7.93 | < .001 | 3.72 | .024 | 8.3 | < .001 |
| 4.07 | .017 | 6.49 | .001 | 3.85 | .021 |
| 4.06 | .017 | 5.36 | .005 | 4.68 | .009 |
| 2.5 | .082 | 5.08 | .006 | 16.54 | < .001 |
| 1.29 | .275 | 0.86 | .422 | 2.18 | .113 |
| 5.73 | .003 | 0.23 | .797 | 12.91 | < .001 |
| | | 6.46 | .001 | 3.52 | .029 |

### Table 10. Bins t-stats with FDR corrected p-values

The following table lists overall, unimodal, and bimodal 2-tailed t-stat values, one for each phase bin (from 1 to 60), with corresponding FDR-adjusted p-values. The yellow filling represents significant (i.e. $p_{FDR} < .05$) t-stat > 0 (slow RTs) while the green filling represents significant (i.e. $p_{FDR} < .05$) t-stat < 0 (fast RTs).

| | overall | unimodal | | bimodal | | | |
|---|---|---|---|---|---|---|---|
| t-stat | p-value (FDR) | t-stat | p-value (FDR) | t-stat | p-value (FDR) | rad | bin |
| 5.14 | < .001 | 2.79 | .036 | 3.38 | .008 | -3.14 | 1 |
| 3.90 | .001 | 2.54 | .051 | 2.95 | .006 | -3.04 | 2 |
| 3.54 | .001 | 2.26 | .052 | 1.63 | .044 | -2.93 | 3 |
| 2.99 | .004 | 2.81 | .041 | 0.16 | .147 | -2.82 | 4 |
| 1.70 | .032 | 2.20 | .049 | -0.22 | 0.141 | -2.72 | 5 |
| 1.37 | .050 | 1.92 | .067 | -0.28 | 0.135 | -2.61 | 6 |
| 1.35 | .051 | 2.01 | .066 | -0.33 | .131 | -2.50 | 7 |
| 0.60 | .118 | 1.56 | .112 | -0.55 | 0.112 | -2.40 | 8 |
| -0.38 | .138 | 0.90 | .239 | -1.10 | 0.074 | -2.29 | 9 |
| -1.48 | .045 | 0.40 | .362 | -1.86 | .031 | -2.18 | 10 |
| -1.93 | .022 | -1.16 | .182 | -1.44 | .048 | -2.08 | 11 |
| -1.98 | .021 | -0.54 | .328 | -1.99 | .026 | -1.97 | 12 |
| -2.05 | .020 | -0.31 | .379 | -2.26 | .02 | -1.86 | 13 |
| -2.88 | .005 | -2.19 | .048 | -2.31 | .019 | -1.76 | 14 |
| -2.82 | .005 | -1.98 | .061 | -1.92 | .029 | -1.65 | 15 |
| -2.77 | .005 | -2.51 | .049 | -1.50 | .047 | -1.54 | 16 |
| -1.37 | .051 | -1.39 | .144 | -0.70 | .109 | -1.44 | 17 |

| | | | | | | | |
|---|---|---|---|---|---|---|---|
| -1.98 | .021 | -1.72 | .085 | -0.67 | .11 | -1.33 | 18 |
| -1.18 | .064 | -1.16 | .177 | -1.07 | .075 | -1.22 | 19 |
| -1.16 | .064 | -1.51 | .120 | -0.52 | .114 | -1.12 | 20 |
| -1.28 | .055 | -1.36 | .146 | -0.74 | .109 | -1.01 | 21 |
| -0.63 | .119 | -0.94 | .235 | -0.56 | .116 | -0.91 | 22 |
| 0.07 | .175 | 0.17 | .420 | -0.89 | .091 | -0.80 | 23 |
| 0.02 | .179 | -0.22 | .409 | -0.34 | .132 | -0.69 | 24 |
| -0.13 | .170 | -0.35 | .371 | -0.03 | .153 | -0.59 | 25 |
| 0.60 | .121 | 0.86 | .245 | -0.67 | .109 | -0.48 | 26 |
| 0.36 | .138 | 0.49 | .341 | 0.40 | .127 | -0.37 | 27 |
| -0.53 | .122 | -0.05 | .433 | 0.11 | .148 | -0.27 | 28 |
| -0.78 | .100 | -0.10 | .421 | -0.64 | .109 | -0.16 | 29 |
| -1.01 | .077 | 0.17 | .413 | -1.34 | .055 | -0.05 | 30 |
| -1.49 | .045 | -0.74 | .269 | -0.56 | .114 | 0.05 | 31 |
| -2.08 | .019 | -0.59 | .316 | -1.22 | .065 | 0.16 | 32 |
| -1.64 | .035 | -1.11 | .187 | -0.89 | .093 | 0.27 | 33 |
| -1.74 | .032 | -1.29 | .159 | -0.96 | .087 | 0.37 | 34 |
| -2.03 | .020 | -1.74 | .088 | -1.20 | .065 | 0.48 | 35 |
| -3.14 | .003 | -2.45 | .051 | -2.51 | .014 | 0.59 | 36 |
| -3.28 | .002 | -2.32 | .052 | -2.36 | .018 | 0.69 | 37 |

| | | | | | | | |
|---|---|---|---|---|---|---|---|
| -2.66 | .006 | -2.32 | .049 | -1.76 | .036 | 0.80 | 38 |
| -2.67 | .006 | -2.21 | .054 | -2.12 | .021 | 0.91 | 39 |
| -2.57 | .007 | -1.98 | .064 | -2.21 | .021 | 1.01 | 40 |
| -2.46 | .009 | -2.00 | .063 | -2.17 | .02 | 1.12 | 41 |
| -2.42 | .010 | -2.35 | .053 | -1.47 | .049 | 1.22 | 42 |
| -2.71 | .006 | -2.40 | .051 | -1.59 | .043 | 1.33 | 43 |
| -1.73 | .031 | -2.20 | .053 | -0.60 | .113 | 1.44 | 44 |
| -1.09 | .071 | -1.23 | .171 | -0.10 | .147 | 1.54 | 45 |
| -0.83 | .096 | -0.75 | .271 | 0.15 | .145 | 1.65 | 46 |
| -0.56 | .119 | -0.90 | .243 | 0.72 | .109 | 1.76 | 47 |
| 0.43 | .133 | -0.42 | .359 | 1.60 | .044 | 1.86 | 48 |
| 0.88 | .091 | 0.01 | .442 | 1.58 | .042 | 1.97 | 48 |
| 0.59 | .117 | -0.11 | .428 | 1.46 | .048 | 2.08 | 50 |
| 1.40 | .050 | 0.85 | .242 | 1.77 | .036 | 2.18 | 51 |
| 2.23 | .014 | 1.22 | .171 | 2.60 | .014 | 2.29 | 52 |
| 3.28 | .002 | 1.84 | .075 | 3.19 | .007 | 2.40 | 53 |
| 3.46 | .002 | 1.73 | .087 | 3.40 | .015 | 2.50 | 54 |
| 3.43 | .002 | 2.10 | .056 | 2.59 | .013 | 2.61 | 55 |
| 4.42 | < .001 | 2.92 | .038 | 2.19 | .02 | 2.72 | 56 |
| 5.18 | < .001 | 3.13 | .029 | 3.12 | .005 | 2.82 | 57 |

| | | | | | | | |
|---|---|---|---|---|---|---|---|
| 5.89 | < .001 | 3.73 | .016 | 3.14 | .005 | 2.93 | 58 |
| 5.57 | < .001 | 3.44 | .019 | 3.16 | .006 | 3.04 | 59 |
| 5.14 | < .001 | 2.79 | .031 | 3.38 | .005 | 3.14 | 60 |

### Linear Mixed Effect Models (LMEMs) for MMSI (magnitude of multisensory integration) and respiration

#### Audio-tactile MMSI

First-base model = MMSI ~ 1 + AUC + (1|participant)

Table 11. model fit stats

| AIC | BIC | Log-Likelihood | deviance |
|---|---|---|---|
| -3455.6 | -3386 | 1740.8 | -3481.6 |

Table 11.a ANOVA marginal tests (on coefficients)

| term | F-stat | DF1 | DF2 | p-value |
|---|---|---|---|---|
| intercept | 1.43 | 1 | 1551 | .232 |
| AUC | 4.02 | 10 | 1551 | < .001 |

Alternative-full model = MMSI ~ 1 + phase + AUC + (1|participant)

Table 12. model fit stats

| AIC | BIC | Log-Likelihood | deviance |
|---|---|---|---|
| -3487.9 | -3402.2 | 1759.9 | -3519.9 |

Table 12.a ANOVA marginal tests (on coefficients)

| term | F-stat | DF1 | DF2 | p-value |
|---|---|---|---|---|
| intercept | 1.10 | 1 | 1548 | .295 |
| AUC | 4.11 | 10 | 1548 | < .001 |
| phase | 12.92 | 3 | 1548 | < .001 |

Table 13. Model comparison: Theoretical Likelihood Ratio Test

| model | DF | AIC | BIC | Log-Likelihood | LR-Stat | delta df | p-value |
|---|---|---|---|---|---|---|---|
| base | 13 | -3455.6 | -3386 | 1740.8 | | | |
| full | 16 | -3487.9 | -3402.2 | 1759.9 | 38.267 | 3 | < .001 |

### Audio-visual MMSI

First-base model = MMSI ~ 1 + AUC + (1|participant)

Table 14. model fit stats

| AIC | BIC | Log-Likelihood | deviance |
|---|---|---|---|
| -2991.3 | -2928.2 | 1507.6 | -3015.3 |

Table 14.a ANOVA marginal tests (on coefficients)

| term | F-stat | DF1 | DF2 | p-value |
|---|---|---|---|---|
| intercept | 0.32 | 1 | 1410 | .572 |
| AUC | 0.86 | 9 | 1410 | .560 |

Alternative-full model = MMSI ~ 1 + phase + AUC + (1|participant)

Table 15. model fit stats

| AIC | BIC | Log-Likelihood | deviance |
|---|---|---|---|
| -3021.8 | -2942.9 | 1525.9 | -3051.8 |

Table 15.a ANOVA marginal tests (on coefficients)

| term | F-stat | DF1 | DF2 | p-value |
|---|---|---|---|---|
| intercept | 2.44 | 1 | 1407 | .118 |
| AUC | 0.88 | 9 | 1407 | .540 |
| phase | 12.32 | 3 | 1407 | < .001 |

Table 16. Model comparison: Theoretical Likelihood Ratio Test

| model | DF | AIC | BIC | Log-Likelihood | LR-Stat | delta df | p-value |
|---|---|---|---|---|---|---|---|
| base | 12 | -2991.3 | -2928.2 | 1507.6 | | | |
| full | 15 | -3021.8 | -2942.9 | 1525.9 | 36.503 | 3 | < .001 |

### Circular Analyses (Response onset)

#### Table 17. Consistency across participants (Response onset)

The following table lists overall, unimodal, and bimodal Z-stats from Rayleigh tests, one for each participant (from 1 to 40), with corresponding p-values. The red filling represents significant (i.e. $p < .05$) non-uniform distribution of the data (response onsets).

| Rayleigh Z | p-value | unimodal | | bimodal | |
|---|---|---|---|---|---|
| 11.05 | < .001 | Rayleigh Z | p-value | Rayleigh Z | p-value |
| 1.54 | .215 | 6.48 | .001 | 5.06 | .006 |
| 10.22 | < .001 | 2.89 | .056 | 0.03 | .974 |
| 0.69 | .502 | 5.37 | .005 | 4.96 | .007 |
| 2.57 | .076 | 1.21 | .297 | 0.05 | .954 |
| 0.62 | .541 | 1.15 | .316 | 1.64 | .193 |
| 16.62 | < .001 | 0.32 | .727 | 0.30 | .742 |
| 2.08 | .125 | 8.97 | < .001 | 7.84 | < .001 |
| 3.83 | .022 | 0.88 | .417 | 2.60 | .075 |
| 0.40 | .671 | 3.33 | .036 | 0.97 | .381 |
| 1.65 | .192 | 0.13 | .875 | 1.21 | .299 |
| 6.57 | .001 | 1.71 | .181 | 0.28 | .758 |
| 2.82 | .059 | 2.57 | .076 | 4.16 | .015 |
| 1.13 | .324 | 2.83 | .059 | 0.63 | .535 |
| 16.69 | < .001 | 1.12 | .326 | 0.86 | .425 |
| 2.07 | .126 | 7.70 | < .001 | 9.95 | < .001 |
| 11.86 | < .001 | 1.40 | .246 | 1.60 | .203 |
| 3.35 | .035 | 7.98 | < .001 | 4.19 | .015 |
| 19.20 | < .001 | 1.67 | .188 | 1.90 | .150 |
| 3.41 | .033 | 12.53 | < .001 | 7.06 | .001 |
| 2.12 | .119 | 1.51 | .221 | 3.84 | .021 |
| 10.21 | < .001 | 2.32 | .098 | 0.57 | .567 |
| 10.14 | < .001 | 4.74 | .009 | 5.50 | .004 |
| 11.17 | < .001 | 6.95 | .001 | 3.51 | .030 |
| 6.30 | .002 | 4.47 | .011 | 6.89 | .001 |
| 2.09 | .124 | 0.10 | .907 | 10.32 | < .001 |
| 0.73 | .480 | 2.93 | .053 | 2.37 | .093 |
| 12.32 | < .001 | 0.65 | .522 | 0.74 | .479 |
| 8.76 | < .001 | 3.65 | .026 | 9.68 | < .001 |
| 4.36 | .013 | 2.93 | .053 | 6.09 | .002 |
| 0.53 | .588 | 4.39 | .012 | 0.93 | .394 |

| | | | | | |
|---|---|---|---|---|---|
| 69.42 | < .001 | 0.22 | .802 | 0.32 | .727 |
| 2.37 | .093 | 41.29 | < .001 | 30.64 | < .001 |
| 4.07 | .017 | 1.22 | .296 | 1.47 | .229 |
| 5.58 | .004 | 0.64 | .529 | 4.22 | .015 |
| 0.57 | .565 | 2.31 | .099 | 3.56 | .028 |
| 13.04 | < .001 | 1.10 | .334 | 0.00 | .995 |
| 10.15 | < .001 | 8.73 | < .001 | 4.71 | .009 |
| 12.49 | < .001 | 9.10 | < .001 | 2.40 | .090 |
| 6.27 | .002 | 11.37 | < .001 | 2.79 | .061 |
| | | 6.11 | .002 | 1.49 | .225 |
